# A highly efficient protocol for transforming *Cuscuta reflexa* based on artificially induced infection sites

**DOI:** 10.1101/2020.04.06.028191

**Authors:** Lena Anna-Maria Lachner, Levon Galstyan, Kirsten Krause

## Abstract

A current bottleneck in the functional analysis of the emerging parasitic model plant *Cuscuta* and the exploitation of its recently sequenced genomes is the lack of efficient transformation tools. Here, we describe the development of a novel highly efficient *Agrobacterium*-mediated transformation protocol for *Cuscuta reflexa* based on the parasitic structure referred to as adhesive disk. Both, *Agrobacterium rhizogenes* and *Agrobacterium tumefaciens* carrying binary transformation vectors with reporter fluorochromes yielded high numbers of transformation events. An overwhelming majority of transformed cells were observed in the cell layer below the adhesive disk’s epidermis, suggesting that these cells are particularly susceptible to infection. Co-transformation of these cells happens frequently when *Agrobacterium* strains carrying different constructs are applied together. Explants containing transformed tissue expressed the fluorescent markers in *in vitro* culture for several weeks, offering a possibility for development of transformed cells into callus.

**ONE SENTENCE SUMMARY:** A protocol that yields high numbers of transformed cells in the adhesive disks of *Cuscuta reflexa* upon exposure to agrobacteria brings closer the vision of generating genetically modified *Cuscuta*.

## INTRODUCTION

The obligate parasitic plant genus *Cuscuta*, commonly known as “dodder”, is found almost worldwide and infects a broad range of host plants. The parasite starts an attack by winding around the host stem. A multicellular feeding structure termed the haustorium, which can reach 2-3 mm in size in some species and originates from a secondary meristem in the cortex close to the vascular bundles, develops within a few days and breaches the host tissue integrity using mechanical pressure and enzymatic digestion of cell walls (Nagar et al., 1984; Johnsen et al., 2015; Kaiser et al., 2015). To make the penetration possible, a sucker-shaped attachment organ provides the necessary counterforce. The development of this organ, termed “adhesive disk” or “upper haustorium” (Vaughn, 2002; Lee, 2007; Kaiser et al., 2015), is easily visible as a swelling of the host-facing side of the parasite’s stem. A bio-adhesive, secreted by the epidermal cells of the adhesive disk, anchors the parasite to the host (Vaughn, 2002; Galloway et al., 2020). Morphologically, the development of the adhesive disk is marked by major local growth processes and shape changes of the involved cells: on one hand elongation of cells in the parasite’s cortex around the developing haustorium takes place; on the other hand, an even more drastic reconfiguration of the epidermis into digitate or club-shaped cells can be noted (Shimizu and Aoki, 2019).

In light of the detrimental effect to important crops exerted by *Cuscuta* worldwide, it is imperative, that mechanisms guiding the parasitic attack are better understood. One prerequisite for achieving a better understanding of the molecular processes and mechanisms of a *Cuscuta* attack is a reference genome. Recently, complete genome sequences have been published for two *Cuscuta* species, *C. campestris* and *C. australis* (Sun et al., 2018; Vogel et al., 2018). However, two significant bottlenecks to perform targeted genome manipulations remain in the form of low transformation efficiency and poor regeneration rates of *Cuscuta* tissues *in vitro*. Some few approaches to transform *C. trifolii* (Borsics et al., 2002) and *C. europaea* (Svubova and Blehova, 2013) have been reported. Explants from seedlings were transformed using *Agrobacterium tumefaciens*, and expression of the transgenes was demonstrated in the calli. However, the generation of a whole transgenic *Cuscuta* plant has to date not been reported, indicating that the current techniques lack efficiency.

Besides *A. tumefaciens, Agrobacterium rhizogenes* has become a popular agent for transforming plant genomes (Ozyigit et al., 2013). *A. rhizogenes* is a soil-borne bacterium infecting many angiosperms and causing them to produce a copious number of roots which became known as the “hairy root syndrome” (Bahramnejad et al., 2019). Like *A. tumefaciens, A. rhizogenes* transfers a segment of DNA known as T-DNA into its hosts. The transfer process is controlled by virulence (*vir*) genes that are induced by phenolic signal molecules (Gelvin, 2003). The *A. rhizogenes* T-DNA is stably integrated into the plant nuclear genome where it expresses the *rol* (*<u>r</u>ooting <u>l</u>ocus*) genes required for excessive adventitious root growth (Ozyigit et al., 2013). What has made these hairy roots popular for plant biotechnology is that they can be propagated in the absence of exogenous plant hormones. Very recently, it was shown that an *A. rhizogenes* gene coding for the mikimopine synthase was horizontally transferred into several *Cuscuta* species (Zhang et al., 2020), including *Cuscuta campestris* (Vogel et al., 2018) and *Cuscuta australis* (Sun et al., 2018), suggesting that *Cuscuta* species may be susceptible to infection by this *Agrobacterium* species despite their lack of roots.

With this study, we set out to test the applicability of the hairy root transformation protocol in *Cuscuta*. Although hairy roots as such were not obtained, we were able to obtain high numbers of transformed cells in the species *Cuscuta reflexa*, particularly in its adhesive disks. We describe here the simple and highly efficient protocol that can yield hundreds of transformed cells within a week based on the use of *A. rhizogenes*. We further show that the protocol is applicable to use with *A. tumefaciens* with equally high success, which widens possibilities for single or co-transformation of different constructs. Our trials with other *Cuscuta* species showed some transformation success but were not as reliable as experiments with *C. reflexa*. Although some species-specific adaptations may be required, our protocol is overall fast, flexible, easily scalable and suitable for transformations of genes at high throughput, and a first step towards making *Cuscuta* amenable to genome modifications.

## RESULTS

### *Agrobacterium rhizogenes* does not induce hairy roots in *Cuscuta* species

*A. rhizogenes* is typically applied to the roots (Ho-Plagaro et al., 2018), hypocotyl (Alagarsamy et al., 2018) or cotyledons (Ron et al., 2014) of dicotyledonous angiosperms by cutting, puncturing or otherwise wounding these tissues. In the absence of proper roots, cotyledons or other leaves, we first tested hairy root induction on young germinating seedlings of *C. campestris* and on different parts of the shoots from different *Cuscuta* species using *A. rhizogenes* with and without the pRedRoot T-DNA. Occasionally, transformed cells exhibiting an intense orange fluorescence from expression of the reporter protein dsRed have indeed been observed, particularly in shoot tips (Supplemental Fig. S1). However, no root development could be observed in any of our trials with the strain MSU440, independent of whether it carried the pRedRoot T-DNA or not and transformation success reliability was poor.

### *A. rhizogenes* transforms adhesive disk cells of *C. reflexa* with high efficiency

We next decided to expose developing infection structures to a pRedRoot-containing *A. rhizogenes* culture. For this we used the method for induction of haustoriogenesis in *C. reflexa* described by Olsen et al. (2016) that uses a combination of far-red light illumination and tactile stimuli to synchronize haustorial development (Tada et al., 1996). The stems on which the infection sites developed where exposed to *A. rhizogenes* for as long as it took for haustoria to emerge (7-8 days) (Fig. 1). Upon microscopical analysis, a large number of intensely orange-fluorescing cells expressing dsRed were revealed that were almost exclusively located in the adhesive disks around the protruding haustorium (Fig. 2A-F). The dsRed fluorescence was visible in distinct spots often consisting of 5-15 clustered cells, but single transformed cells and bigger clusters were also observed (Fig. 2). Cross sections through sites where transformation had occurred revealed that the cells expressing the dsRed were mostly not located at the very surface but rather in a cell layer directly below the elongated epidermal cells (Fig. 2G-L). 52 % of the adhesive disks exhibited one or more spots with dsRed fluorescence (based on 52 infected shoots with 426 infection structures) (Table 1), but there was considerable variation between individual shoots. While some green and blue autofluorescence was observed in the central haustorial tissue (Supplemental Fig. S2), the adhesive disk of *C. reflexa* exhibited little to no autofluorescence, as demonstrated by mock transformations with *A. rhizogenes* cells that lack the pRedRoot T-DNA (Fig. 2M, N). Experiments where *A. rhizogenes* was removed or added after 2 days, showed that the uptake of the T-DNA in the first two days is minimal to absent, and seems to happen only once the development of the infection sites has commenced (Supplemental Tab. S1, 2).

**Table 1:**
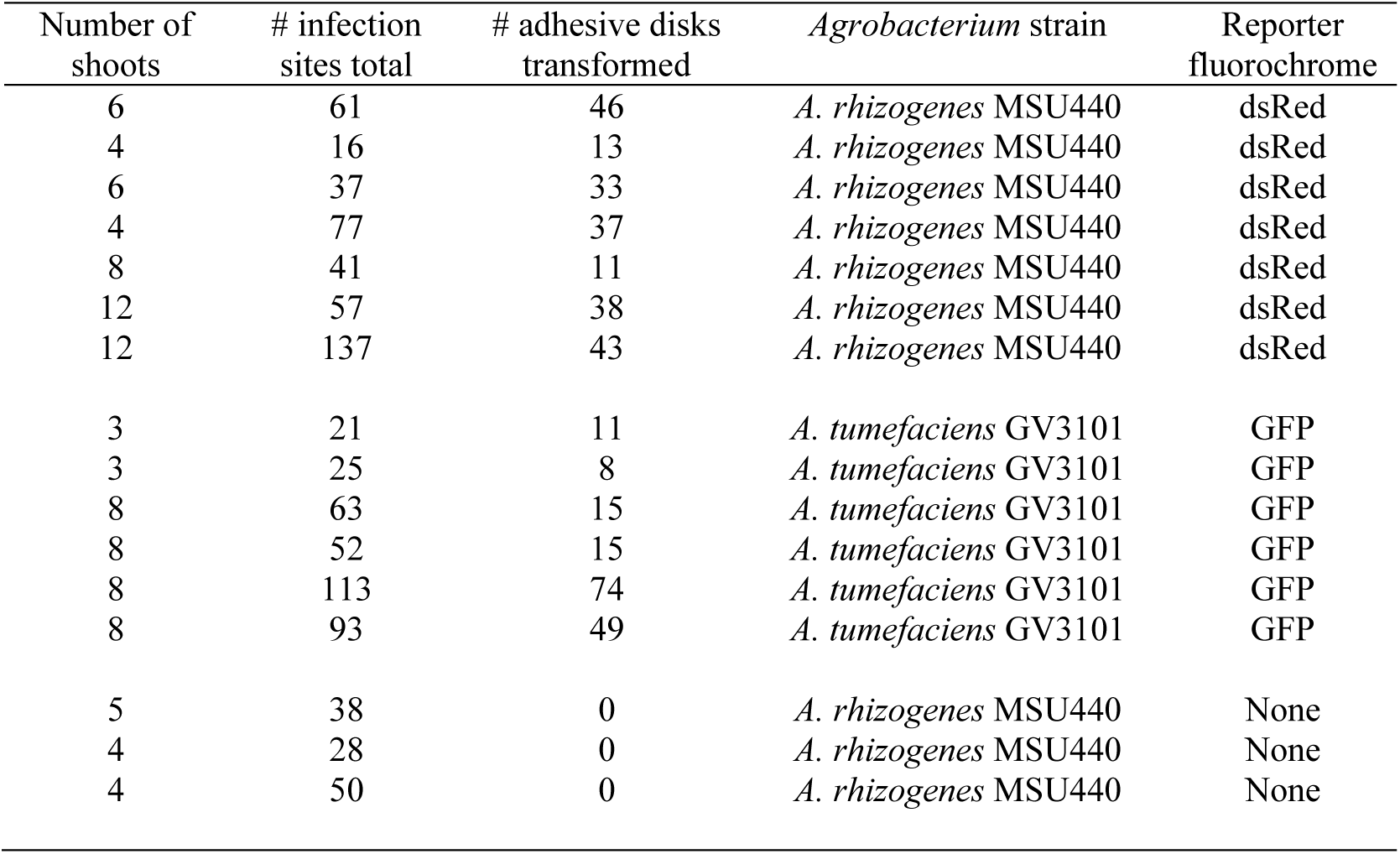
Transformation efficiency overview

**Figure 1:**
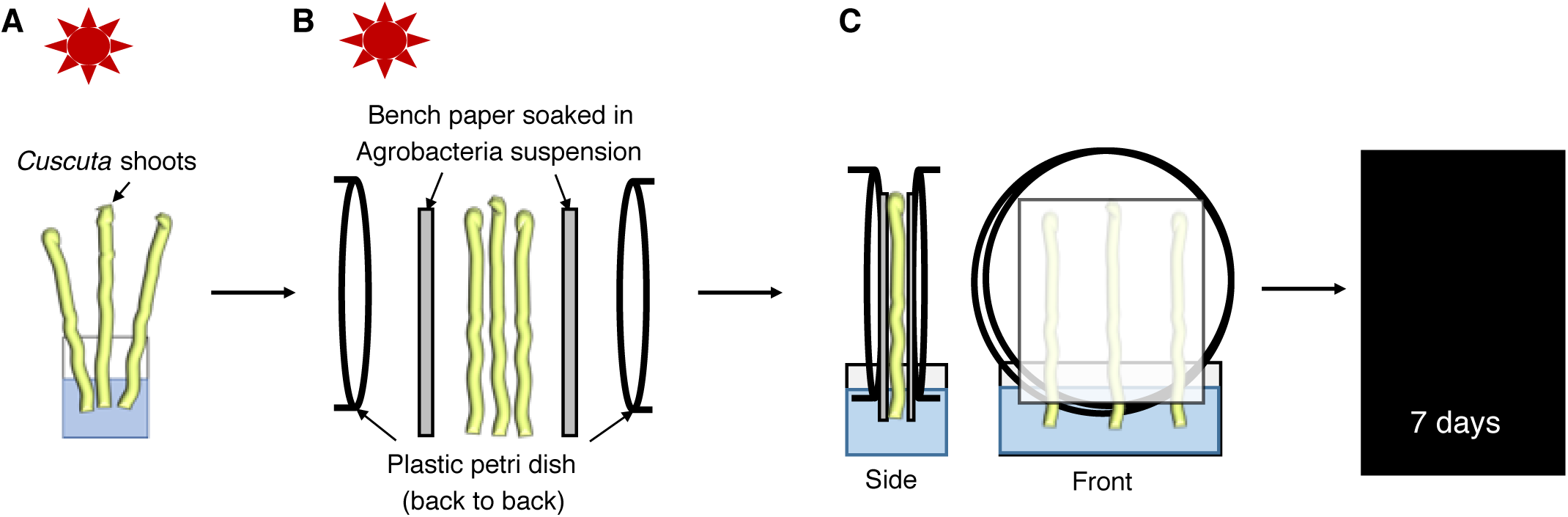
Overview of the experimental setup. A, Exposure of *Cuscuta* shoots to far red light (740 nm) for 90-120 min. B, Placement of *Cuscuta* shoots between *Agrobacterium*-soaked bench paper sheets and two petri dish halves (back to back). C, Incubation in darkness with shoots and bench paper placed in a water reservoir.

**Figure 2:**
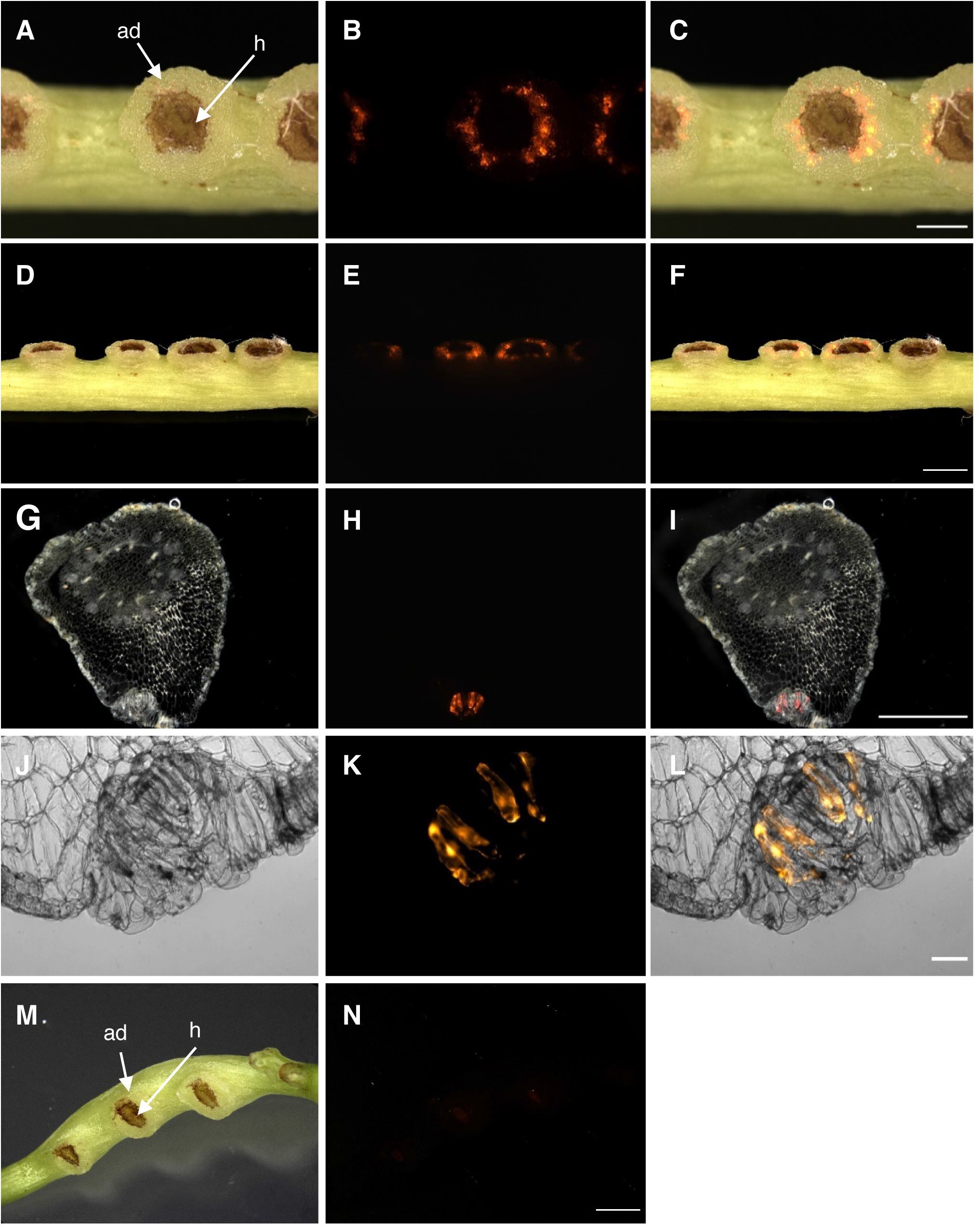
Transformation of *C. reflexa* adhesive disks by *A. rhizogenes* containing the binary vector pRedRoot. A-F, Intact infection sites after transformation. Topview (A-C) and sideview (D-F) of transformed adhesive disks are shown. G-L, Semithin vibratome sections of transformed adhesive disk tissue showing sub-epidermal localization of transformed cells. M-N, Mock transformation with *A. rhizogenes* lacking the binary pRedRoot plasmid. Darkfield or brightfield pictures (first column) are shown alongside the fluorescence images taken with a Cy3 filter (middle column). Overlays of both are shown in the right column. Adhesive disks (ad) and haustoria (h) are indicated. Scale bars are 1000 µm (C, I), 2000 µm (F, N) and 100 µm (L).

### Adhesive disks readily take up the live cell stain 5-carboxyfluorescein-diacetate (CFDA)

Life cell stains like CFDA are membrane permeable and are hydrolyzed in the cytoplasm to the green fluorescent carboxyfluorescein. Fluorescence inside cells is thus on one hand evidence for their viability and on the other hand reveals which cells lack protection by barriers such as the cuticle that would block uptake of the stain. When shoots that showed dsRed fluorescence in the adhesive disks where exposed to a CFDA solution, the adhesive disks and often (but not always) also the haustoria exhibited green fluorescence (Fig. 3). CFDA fluorescence was also observed regularly in young side-shoot buds but only very rarely in the intact stems (Supplemental Fig. S1). This indicates that the cuticle that protects *Cuscuta* stems and that hinders the uptake of the stain into stems is most likely “leaky” or absent in the adhesive disks and haustoria, thus permitting the stain to enter the respective tissue. This, in turn may explain its susceptibility to agrobacterial infection.

**Figure 3:**
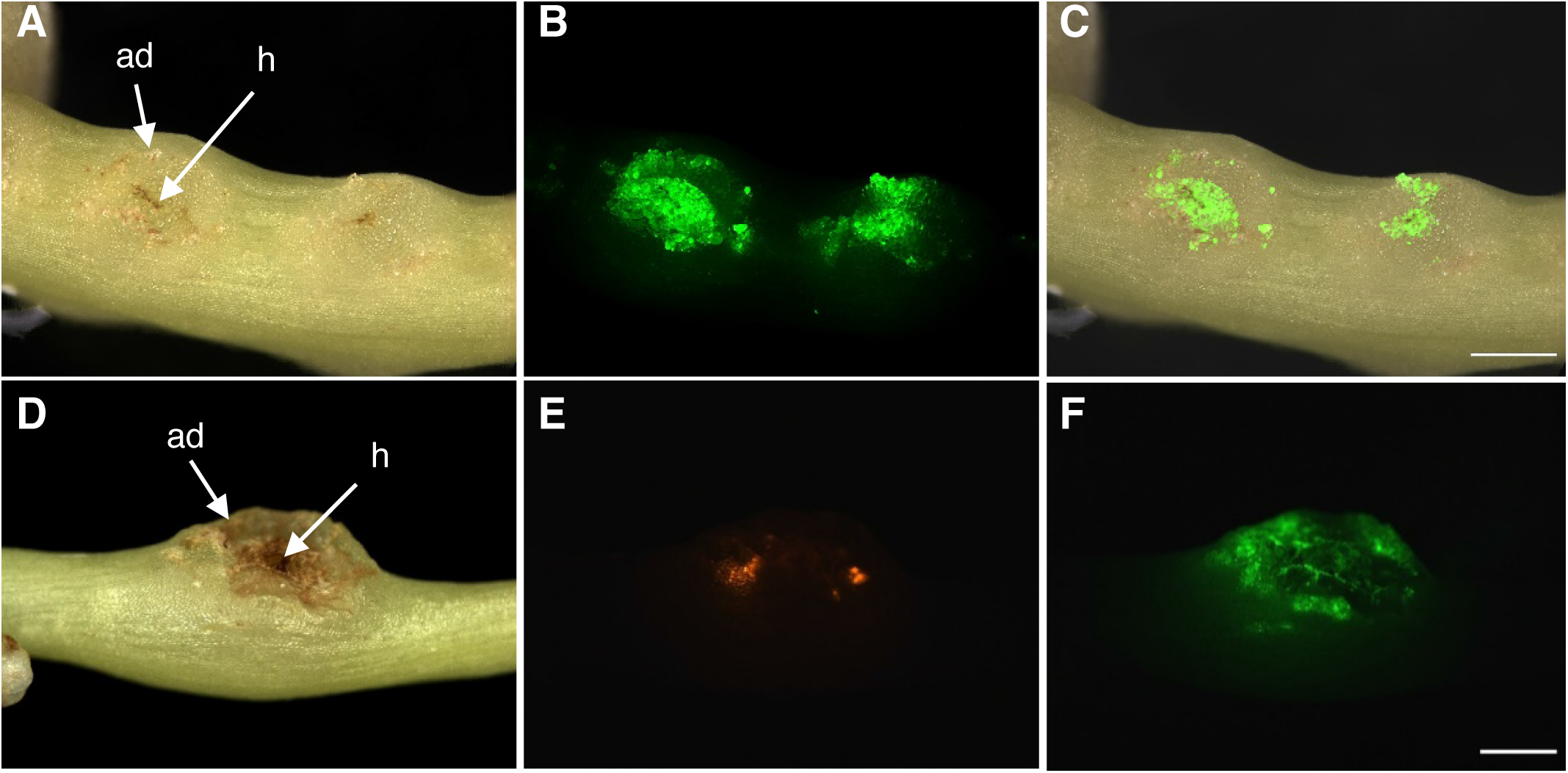
Uptake of 5-Carboxyfluorescein-diacetate (CFDA) into adhesive disks of *C. reflexa*. A-C, Green CFDA fluorescence is shown in a shoot segment containing two developing infection sites. A superimposition of the images from A and B is shown in C. D-F, CFDA uptake into an adhesive disk containing dsRed-expressing cells. Darkfield images (left column) and fluorescence images (middle column and lower right image) are shown. Adhesive disks (ad) and haustoria (h) are indicated. Scale bars are 1000 µm.

### Application of the transformation protocol is not limited to *A. rhizogenes*

In order to reveal whether the high transformation rates were a result of a specific susceptibility of *C. reflexa* to *A. rhizogenes* we exposed far-red light induced stems to *Agrobacterium tumefaciens* carrying a GFP gene in the T-DNA of a binary plasmid (Bobik et al., 2019) using the same setup. Only very weak background fluorescence was seen in this case in the orange channel and in the blue channel, while the high intensity of green fluorescence in a ring corresponding to the adhesive disk indicated that the GFP was expressed in this tissue as a result of the transformation (Fig. 4A-I). As with the dsRed, GFP was expressed in elongated cells beneath the layer of epidermal cells in the adhesive disk (Supplemental Fig. S3). The infection frequency was on average 43% for *A. tumefaciens* (based on 38 infected shoots with 367 infection structures) (Tab. 1).

**Figure 4:**
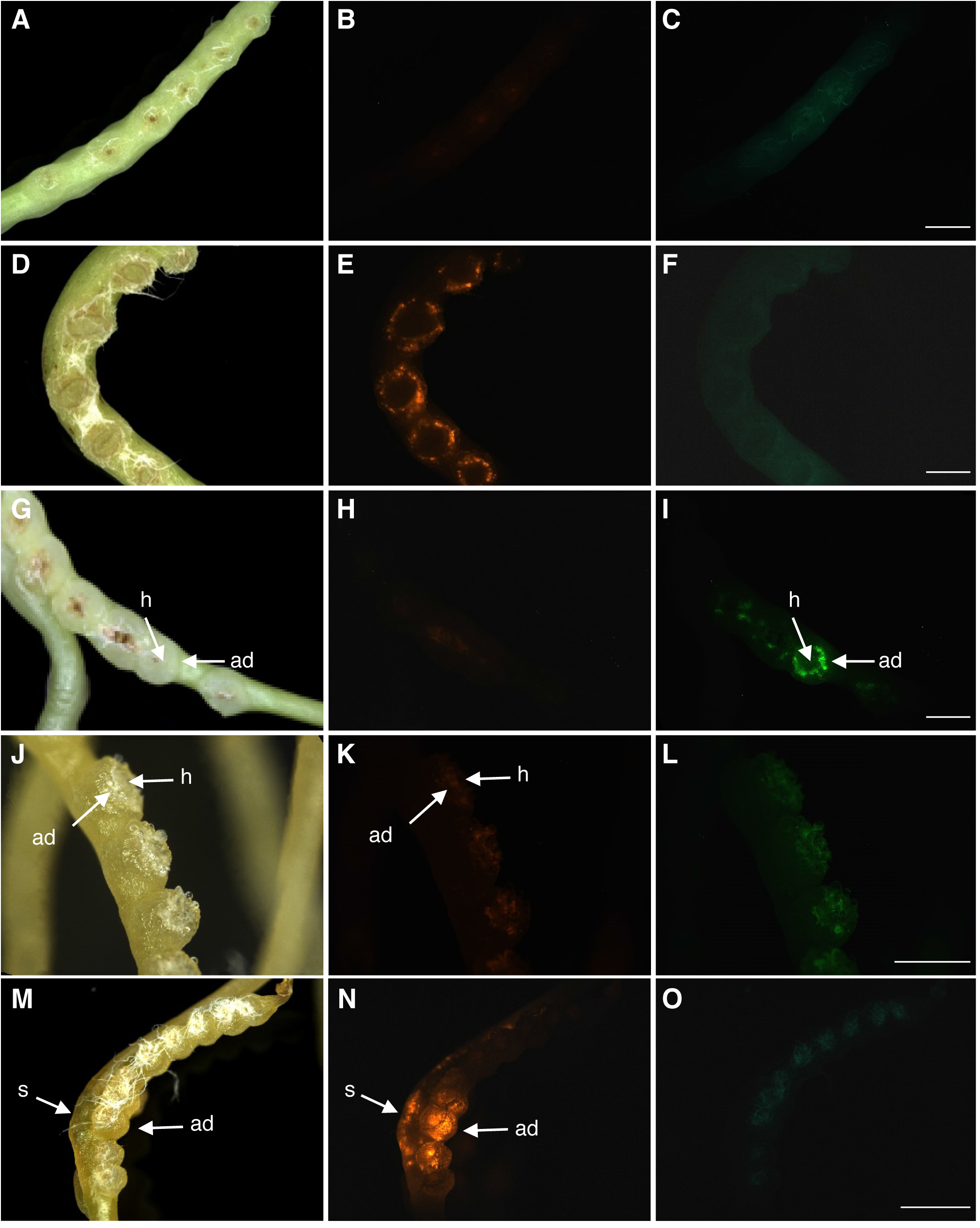
Extension of the protocol to *A. tumefaciens* and *C. campestris*. A-F, Negative (A-C) and positive (D-F) controls using the combination of *C. reflexa* and *A. rhizogenes* (see also Fig. 2). G-I, Transformation after combining *C. reflexa* with *A. tumefaciens* containing a binary GFP-expressing vector. J-O, Negative control (J-L) and pRedRoot transformation (M-O) using the combination of *C. campestris* and *A. rhizogenes*. Scalebars represent 2000 µm (C, F, I, O) and 1000 µm (L), respectively. White fibres of the bench paper from the experimental setup can be seen adhering strongly to the adhesive disks in some darkfield images (D, G, J, M).

### Transformation of other *Cuscuta* species

Within the genus *Cuscuta*, three subgenera are distinguished: subgenus *Monogyna*, which includes *C. reflexa*, subgenus *Grammica* and subgenus *Cuscuta* (Yuncker, 1932). To test whether our protocol is applicable to *C. campestris* whose sequenced genome (Vogel et al., 2018) would make it a very interesting target for genome modifications we repeated the same transformation setup with this species (Fig. 4) and a third *Cuscuta* species, *C. platyloba* (Supplemental Fig. S1), both belonging to the subgenus *Grammica*. With both *Agrobacterium* species, a higher degree of necrotic tissue was observed in these two species as a result of this treatment, which, in turn, created a higher amount of unspecific autofluorescence. While adhesive disk transformation could be observed in *C. campestris* (Fig. 4K-L), it was by far not as frequent as in *C. reflexa* and was often weaker than in stem tissue adjacent to haustoria-forming sites.

### Simultaneous exposure to both *Agrobacterium* strains yields a high number of co-transformation events

A desired feature of transformation protocols is the possibility to express multiple transgenes in the same cell. This can be achieved by time-consuming sequential transformation or the cloning of suites of genes into the T-DNA of one vector, often yielding large unwieldy constructs. However, the high susceptibility of the same *C. reflexa* tissue to both, *A. rhizogenes* and *A. tumefaciens*, opens for the possibility of introducing multiple constructs into the same cell by co-infection. To achieve this, both species of *Agrobacterium* carrying each their respective fluorescent reporter construct (dsRed in *A. rhizogenes* and GFP in *A. tumefaciens*) were mixed in a 1:1 ratio (based on their ODs at 595 nm) prior to exposing the *C. reflexa* stems to them in our transformation setup. Fluorescence microscopy revealed that both, dsRed and GFP were visible with similar yields in the adhesive disks as in single transformation experiments. A considerable amount of overlapping fluorescence indicated that co-transformation did in fact occur at a high rate (Fig. 5A-D). In order to see whether the same cells (and not just cells in the same area) indeed expressed both fluorescent proteins, we prepared semi-thin cross sections through transformed regions and documented the fluorescence location with microphotography (Fig. 5E-H) and by densitometry scanning of fluorescence intensities over an area containing several transformed clusters (Fig. 5I, Supplemental Fig. S4). Both revealed an exact coincidence of the two fluorophores in several cells, suggesting that there are hot spots of susceptible tissue that is frequently co-transformed. At the same time, the occurrence of cells transformed with only one fluorochrome shows that each fluorescence signal is specific.

**Figure 5:**
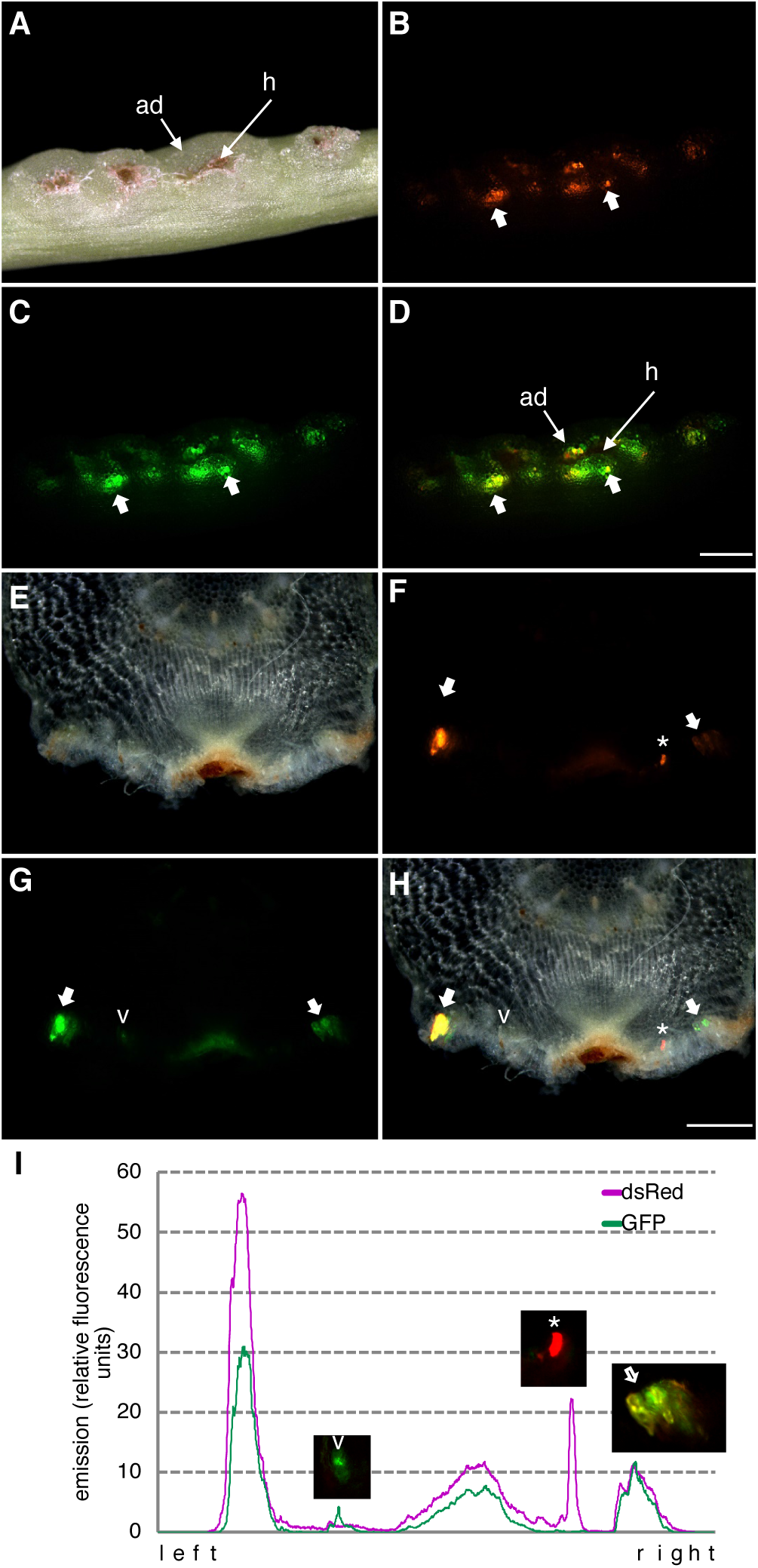
Co-transformation of different reporter proteins in *C. reflexa* adhesive disks. DsRed was introduced with *A. rhizogenes* while GFP was introduced with *A. tumefaciens*. A-D, Side view of intact stem with darkfield (A), red fluorescence (B) and green fluorescence (C) images. D shows a superimposed image of the two fluorescence images with thick arrows pointing at spots where both reporter proteins coincide (yellow color due to overlay). The scalebar measures 1000 µm. E-H, Cross section through a transformed infection site with darkfield (E), red fluorescence (F) and green fluorescence (G) images. H shows a superimposed image of all three images. The scalebar measures 500 µm. The asterisk and the arrowhead indicate cells that are transformed with only one reporter protein. The thick arrow indicates cells were both reporter proteins coincide. I, Intensity scan performed on the two single fluorescent images with the purple line representing the dsRed fluorescence and the green line representing the GFP fluorescence.

### Transgene expression after a transformation event is preserved over several weeks

The frequent occurrence of transformed cell clusters raised the question whether these arose through cell division and propagation of single transformed cells, indicating not only a stable insertion of the transgenes but, moreover, the possibility to regenerate vegetative or reproductive tissue by *in vitro* propagation from the transformation events. To test this, we sterilized explants containing transformed tissue and maintained them in *in vitro* cultures. The explants showed slight growth of cells at the edges of the adhesive disks, including the transformed regions, but significant propagation was not observed over a period of up to 8 weeks. The fluorescence was consistently high for at least 4 weeks (Fig. 6) but started to decreased during longer cultivation times.

**Figure 6:**
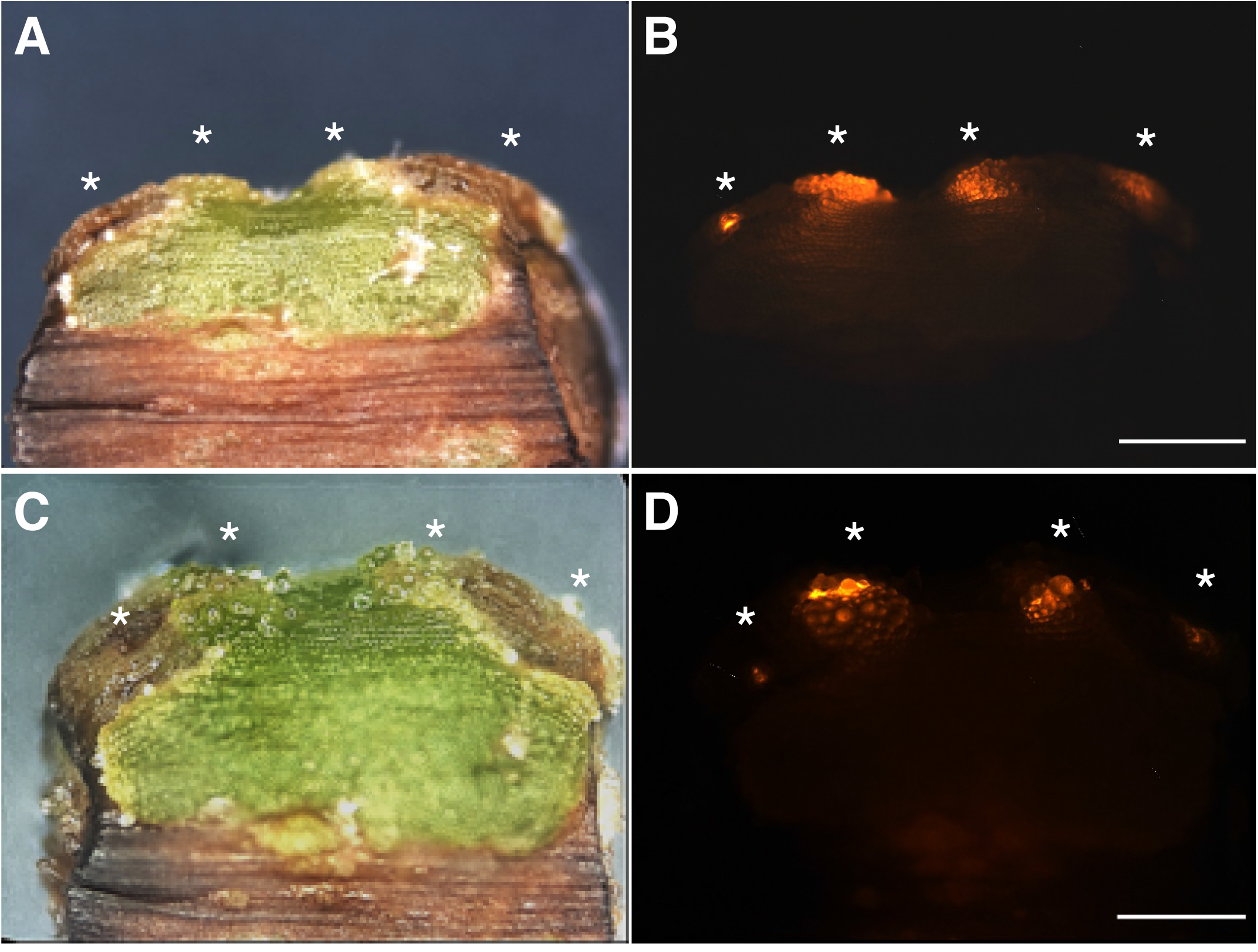
Cultivation of pRedRoot-expressing explants from *C. reflexa*. The shown explant contains two infection structures with transformed adhesive disks. Asterisks indicate clusters of transformed cells. Images were taken after 5 days (A-B) and 4 weeks (C-D) in sterile culture. Brightfield images are shown next to the red fluorescence. The scalebars represent 1000 µm.

## DISCUSSION

In the last few years, research on *Cuscuta* has seen a rapid increase and with the publication of the first two genome sequences of *C. campestris* (Vogel et al., 2018) and *C. australis* (Sun et al., 2018), our knowledge on these parasites has experienced a significant leap forward. Nevertheless, there are still several obstacles that need to be overcome before the possibilities offered by these genomes can be fully exploited. The main bottleneck is the lack of an efficient and reproducible protocol with which targeted genetic manipulations can be performed. Previous attempts to transform *Cuscuta* have shown that *A. tumefaciens* is able to transform callus cells (Borsics et al., 2002; Svubova and Blehova, 2013). However, in our hands these transformation events were very scarce, explaining maybe why this approach has not yielded greater success.

With the work presented here, we show that transformation of *Cuscuta reflexa* works very efficiently when developing infection sites are targeted. As our experiments show, the protocol works with both, *A. rhizogenes* and *A. tumefaciens* and allows the expression of reporter proteins from different binary vectors and under the control of different promoters. The T-DNA of pRedRoot, a binary vector developed for *A. rhizogenes*, contains the dsRed protein under control of the Ubi10 promoter (Libiakova et al., 2018), while the *A. tumefaciens* line used by Bobik et al. (2019) contains a GFP gene controlled by the 35S promoter. Both promoters allow strong constitutive expression of transgenes. It remains to be shown if other - in particular inducible - promoters work in *Cuscuta* as well as the standard constitutive promoters do, or if adaptations are required for use with the parasite.

The simplicity of the method (low-tech, cheap and easy-to-learn) (Fig. 1) and the high reproducibility makes it suitable for larger-scale screening approaches. The high numbers of transformed cells and the longevity of transgene expression are good preconditions for the development of a callus culture that can then be used to regenerate transgenic *Cuscuta* plants.

The high transformation rates of adhesive disk cells is certainly noteworthy. Agrobacteria are known to be attracted to polyphenolic substances exuded by plant roots (Ozyigit et al., 2013). Haustorial sites are also rich in hydroxycinnamic acid derivatives, caffeic acid depsides and other polyphenols (Löffler et al., 1995), which is evident by the rapid browning of infection sites upon their exposure to air (Johnsen et al., 2015). Comparably high concentrations of phenolic substances, albeit in a slightly different composition, are interestingly also found in the meristematic apex of *Cuscuta* shoots (Löffler et al., 1995). After adhesive disks, shoot tips were the second tissue that showed a heightened susceptibility towards agrobacterial infection. However, attraction by polyphenols does not explain why only the adhesive disks and not the haustoria are transformed as the latter exhibit high phenolic substance production as well (Löffler et al., 1995). Both haustoria and the surrounding adhesive disks are characterized by a high metabolic activity, so that limiting rates of transcription and translation are not likely to be the cause for the lack of transgene expression in the haustorium. The life cell stain CFDA that indicates accessibility and viability of stained cells was also detected in both organs (Fig. 3), albeit with a preference for adhesive disks over haustoria. Stems, in contrast were not stained. The presence (in stems) or absence (in infection sites) of a cuticle supposedly determines how rapidly the stain is absorbed. The cuticle could likewise be effective in reducing access to cells for agrobacteria. However, even with these explanations, it remains a mystery why the adhesive disks show high transformation rates while the haustoria are not transformed at all, and this certainly deserves further investigations in the future.

None of the species that belong to the genus *Cuscuta* possesses roots and they are therefore no obvious natural targets of the plant pathogenic bacterium *Agrobacterium rhizogenes*. Nevertheless, some *Cuscuta* species were recently shown to have acquired a gene coding for the Mikimopine synthase (*mis* gene) (Zhang et al., 2020) that is typically transferred to plants during *A. rhizogenes* infection in order to supply the bacterium with opines. Plant homologues of the *mis* gene are found only in a handful of plant species belonging to the genera *Nicotiana* and *Linnaria* where they are believed to have originated by three independent horizontal gene transfer (HGT) events (Kovacova et al., 2014). These species do not display the hairy root syndrome, which hypothetically could be attributed to the HGT-derived *mis* gene. By way of small interfering RNAs from the HGT-derived *mis* that may degrade T-DNA-borne *mis* transcripts during an infection, these acquired genes could potentially prevent *A. rhizogenes* growth. Their evolution under selective pressure and the coverage of *mis*-derived siRNAs, at least, seem to corroborate this possibility for *Nicotiana* (Kovacova et al., 2014). It can be debated whether this could also be the case in *Cuscuta*. However, the *mis* gene was so far only found in species belonging to the subgenus *Grammica* (Zhang et al., 2020), but was not detected in a transcriptome database (Olsen et al., 2016) of *C. reflexa*. Therefore, it is more likely that the loss of key genes involved in root development is responsible for the failure to produce hairy roots in *A. rhizogenes*-infected *Cuscuta* tissue.

## CONCLUSIONS

Using fluorescent reporter proteins and far-red light mediated infection structure induction, we have shown that the adhesive disk of *C. reflexa* is highly susceptible to *Agrobacterium*-mediated transformation. With the high number of transformation events that were observed using our protocol and with the stability of transgene expression, it will be possible to perform transformations with a high number of constructs. Applications of this protocol are, among others, protein localization studies, protein interaction studies (using the high co-transformation rates) and expression of heterologous or synthetic transgenes. Moreover, if the transformed cells can be induced to produce callus and ultimately whole regenerated plants, it will finally be possible to harness the genome sequence information and create *Cuscuta* mutants.

## MATERIALS AND METHODS

### Plant material and *Agrobacterium* strains

*Cuscuta reflexa, Cuscuta campestris* and *Cuscuta platyloba* were grown in a greenhouse on the host *Pelargonium zonale* under continuous illumination and a constant temperature of 21 °C (Förste et al., 2019). The bacteria and binary T-DNA-containing vectors pRedRoot and XM82, respectively, were kindly contributed by Prof. Harro Bouwmeester (University of Amsterdam, Netherlands) (*A. rhizogenes*) and Prof. Tessa Burch-Smith (University of Tennessee, USA) (*A. tumefaciens*) and are described in more detail in other studies (Limpens et al., 2004; Libiakova et al., 2018; Bobik et al., 2019). Cultures of *A. rhizogenes* MSU440 and *A. tumefaciens* GV3101 without binary plasmids were grown on LB medium (tryptone 10 g/L, NaCl 10 g/L, yeast extract 5 g/L, agar 7,5 g/L) supplemented with 100 mg/L Spectinomycin or 50 mg/L Rifampicin plus 50 mg/L Gentamycin, respectively. For bacteria containing the respective binary vectors, Kanamycin at 50 µg/ml was added to the growth medium.

### Induction of infection structure formation by far-red light

For induction of infection structures, *Cuscuta* apical shoots of approximately 12 cm were harvested from the stock plant and exposed to far red light (740 nm) in an otherwise dark room for 90-120 minutes as described before (Olsen et al., 2016) with modifications. The steps were conducted in far red light to not reverse the induction (Tada et al., 1996). To provide a tactile stimulus, four shoots of roughly equal diameter were placed next to each other between two layers of bench paper with one-sided plastic coating (Whatman® Benchkote® surface protector; Cat. # 2300731) that was moistened with tap water (filter-paper side facing the shoots). This set-up was carefully placed between two back-to-back facing petri dish halves (with the filter paper and the cut ends both protruding into a container with tap water (see Fig. 1). Moderate pressure was applied by taping the two petri dish plates together. When kept at room temperature in darkness, infection structures started to develop after about three days (Olsen et al., 2016).

### Transformation of *Cuscuta* cells

An *Agrobacterium* culture was grown overnight in selective media (see above) and adjusted to an OD (595 nm) of 1 to 1.6, before using 2 ml of this suspension to soak the paper side of the bench paper (approximately 8×8 cm area). For mock controls, agrobacteria lacking a T-DNA-encoding binary vector were used. The assembly with far red light-treated *Cuscuta* stems was done as described above and the whole setup was then incubated in a dark incubator set to room temperature for seven days. After disassembling the setup, shoots were briefly rinsed under tap water, remnants of filter paper sticking to the adhesive disks were carefully removed without damaging the plant tissue and stems were kept for up to two days in water or wrapped in wet paper towels before being subjected to microscopical analysis.

For exchange of water to *Agrobacterium* culture and *vice versa* (see Supplemental tables), the setup was disassembled on day 3 under far red light (740 nm) and the bench paper layer exchanged before the setup was re-assembled and subjected to further incubation in the dark for another five days.

### Microscopical imaging

Fluorescence localization in the *Cuscuta* stems exposed to agrobacteria and in the corresponding controls was documented using a StereoLumar V12 stereo microscope or an Axiovert M200 inverted microscope (both from Zeiss) using Zeiss filter sets 43 (for dsRed) and 38 (for GFP). Images were taken using the Axiovision Software (Version 4.8.2). The same exposure times were used for the different fluorescence filter sets for a given sample or magnification, unless specified otherwise. FIJI/ImageJ (V 2.0.0) was used to analyze the pictures, add scalebars, and produce overlays. When adjusting brightness, contrast, minimum and maximum intensities, all fluorescence images of one set were treated alike.

Fluorescence intensity scanning was performed on the marked area containing the region of interest (see Supplemental Fig. S4) using the histogram function of FIJI/ImageJ. Intensity counts were exported to Microsoft Excel for visualization in one joint colored graph.

### Vibratome sectioning

Transformed infection sites were cross-sectioned using a Vibratome (Leica VT1000 E vibrating blade microtome). Section thickness was 100 µm. Sections were viewed and documented using a StereoLumar V12 stereo microscope or an Axiovert M200 inverted microscope (Zeiss) using Zeiss filter sets 43 (for dsRed) and 38 (for GFP).

### Life cell staining with 5-carboxyfluorescein di-acetate (CFDA)

To evaluate the vitality of the transformed cells, the vital stain CFDA (50 mM in DMSO) was diluted immediately prior to use to a final working concentration of 1 mM in water. Stems were covered with a thin layer of CFDA by spreading small drops of a few µl each evenly over the *Cuscuta* stem and infection sites (adjusted from “Drop-And-See assay” (Cui et al., 2015)). After incubation for 10 minutes in the dark, the stems were rinsed with tap water, gently dried with paper and viewed under a StereoLumar V12 stereo microscope using Zeiss filter set 38.

### Cultivation of explants

*Cuscuta reflexa* stems with infection sites and confirmed transformation events were sterilized for 2-5 minutes in 70 % Ethanol and during this time gently cleaned using a brush. After a subsequent 15 minutes incubation step in 1.2 % Sodiumhypochloride, the stems were incubated on a shaker in sterile tap water containing 400 mg/L Cefotaxime overnight, then cut into pieces that contained one or two infection structures each and transferred to MS (Murashige and Skoog) Basal Medium supplemented with 0.8 % Agar, 3 % Sucrose, MS Vitamin solution and 400 mg/L Cefotaxime. Plates were covered with aluminum foil to avoid photobleaching and were kept at 23 °C. Explants were transferred to fresh medium approximately every 4 weeks.

## ACKNOWLEDGEMENTS

This work is part of the doctoral thesis of LL. Financial support from the Tromsø Research Foundation (Mohn Foundation) (grant 16-TF-KK to KK) and the Department of Arctic and Marine Biology (AMB, UiT The Arctic University of Norway) is gratefully acknowledged. We thank Prof. Harro Bouwmeester (University of Amsterdam, Netherlands) for stimulating discussions that provided the initial impulse to this study. We further thank the “Norway-Armenia cooperation in plant molecular biology and biotechnology for agricultural development” project (CPEA-LT-2016/10092), funded by the Eurasia program, Norwegian Agency for International Cooperation and Quality Enhancement in Higher Education (DIKU) for supporting the research visit of LG to Tromsø. Staff at the Climate Laboratory Holt (especially Leidulf Lund) is thanked for *Cuscuta* maintenance. *Agrobacterium* strains and binary reporter plasmids were kindly provided by Prof. Tessa Burch-Smith (University of Tennessee, USA) and Prof. Harro Bouwmeester (University of Amsterdam, Netherlands). Prof. Karsten Fischer and Dr. Stian Olsen (both from UiT The Arctic University of Tromsø, Norway) have contributed with numerous suggestions and discussions to the work and the manuscript.

## SUPPLEMENTARY INFORMATION

The following supplemental materials are available:

**Supplemental Table 1:** Transformation efficiency in *C. reflexa* shoots when *Agrobacterium* solution was replaced with water on day 3 of infection structure induction.

**Supplemental Table 2:** Transformation efficiency in *C. reflexa* shoots when *Agrobacterium* solution was added on day 3 of infection structure induction.

**Supplemental Figure S1**. Examples for infrequent events of transformation in shoot tip and stem.

**Supplemental Figure S2**. Specific fluorescence and autofluorescence of an adhesive disk transformed with *A. rhizogenes* carrying the binary plasmid pRedRoot.

**Supplemental Figure S3**. Localization of GFP expressing cells in semi-thin adhesive disk sections of *C. reflexa* upon transformation with *A. tumefaciens*.

**Supplemental Figure S4**. Area used for intensity-scanning of the images taken of a co-transformed adhesive disk (related to Fig. 5I).

## Notes

Funding information: The work was funded by the Tromsø Research Foundation (grant 16-TF-KK to K. K.), UiT The Arctic University of Norway (PhD stipend to L. L.) and the Norwegian Agency for International Cooperation and Quality Enhancement in Higher Education (DIKU) (Eurasia program grant CPEA-LT-2016/10092, supporting a research visit by L. G. to Tromsø).

## LITERATURE CITED

Alagarsamy K, Shamala LF, Wei S (2018) Protocol: high-efficiency in-planta *Agrobacterium-*mediated transgenic hairy root induction of *Camellia sinensis var. sinensis*. Plant Meth 14: 17

Bahramnejad B, Naji M, Bose R, Jha S (2019) A critical review on use of *Agrobacterium rhizogenes* and their associated binary vectors for plant transformation. Biotechnol Adv 37: 107405

Bobik K, Fernandez JC, Hardin SR, Ernest B, Ganusova EE, Staton ME, Burch-Smith TM (2019) The essential chloroplast ribosomal protein uL15c interacts with the chloroplast RNA helicase ISE2 and affects intercellular trafficking through plasmodesmata. New Phytol 221: 850–865

Borsics T, Mihalka V, Oreifig AS, Barany I, Lados M, Nagy I, Jenes B, Toldi O (2002) Methods for genetic transformation of the parasitic weed dodder (*Cuscuta trifolii* Bab. et Gibs) and for PCR-based detection of early transformation events. Plant Sci 162: 193–199

Cui W, Wang X, Lee JY (2015) Drop-ANd-See: a simple, real-time, and noninvasive technique for assaying plasmodesmal permeability. Methods Mol Biol 1217: 149–156

Förste F, Mantouvalou I, Kanngiesser B, Stosnach H, Lachner LA, Fischer K, Krause K (2019) Selective mineral transport barriers at *Cuscuta*-host infection sites. Physiol Plant doi: 10.1111/ppl.13035

Galloway AF, Knox P, Krause K (2020) Sticky mucilages and exudates of plants: putative microenvironmental design elements with biotechnological value. New Phytol 225: 1461–1469

Gelvin SB (2003) *Agrobacterium*-mediated plant transformation: the biology behind the “gene-jockeying” tool. Microbiol Mol Biol Rev 67: 16–37

Ho-Plagaro T, Huertas R, Tamayo-Navarrete MI, Ocampo JA, Garcia-Garrido JM (2018) An improved method for *Agrobacterium rhizogenes*-mediated transformation of tomato suitable for the study of arbuscular mycorrhizal symbiosis. Plant Meth 14: 34

Johnsen HR, Striberny B, Olsen S, Vidal-Melgosa S, Fangel JU, Willats WG, Rose JK, Krause K (2015) Cell wall composition profiling of parasitic giant dodder (*Cuscuta reflexa*) and its hosts: a priori differences and induced changes. New Phytologist 207: 805–816

Kaiser B, Vogg G, Furst UB, Albert M (2015) Parasitic plants of the genus *Cuscuta* and their interaction with susceptible and resistant host plants. Front Plant Sci 6: 45

Kovacova V, Zluvova J, Janousek B, Talianova M, Vyskot B (2014) The evolutionary fate of the horizontally transferred agrobacterial mikimopine synthase gene in the genera *Nicotiana* and *Linaria*. PLoS One 9: e113872

Lee KB (2007) Structure and development of the upper haustorium in the parasitic flowering plant *Cuscuta japonica* (Convolvulaceae). Am J Bot 94: 737–745

Libiakova D, Ruyter-Spira C, Bouwmeester HJ, Matusova R (2018) *Agrobacterium rhizogenes* transformed calli of the holoparasitic plant *Phelipanche ramosa* maintain parasitic competence. Plant Cell Tiss Organ Cult 135: 321–329

Limpens E, Ramos J, Franken C, Raz V, Compaan B, Franssen H, Bisseling T, Geurts R (2004) RNA interference in *Agrobacterium rhizogenes*-transformed roots of *Arabidopsis* and *Medicago truncatula*. J Exp Bot 55: 983–992

Löffler C, Sahm A, Wray V, Czygan F-C, Proksch P (1995) Soluble phenolic constituents from *Cuscuta reflexa* and *Cuscuta platyloba*. Biochem Syst Ecol 23: 121.128

Nagar R, Singh M, Sanwal GG (1984) Cell wall degrading enzymes in *Cuscuta reflexa* and its hosts. J Exp Bot 35: 8

Olsen S, Striberny B, Hollmann J, Schwacke R, Popper Z, Krause K (2016) Getting ready for host invasion: elevated expression and action of xyloglucan endotransglucosylases / hydrolases in developing haustoria of the holoparasitic angiosperm *Cuscuta*. J Exp Bot 67: 695–708

Ozyigit II, Dogan I, Tarhan EA (2013) *Agrobacterium rhizogenes*-mediated transformation and its biotechnological applications in crops. In KR Hakeem, P Ahmad, M Ozturk, eds, Crop Improvement. Springer Science + Business Media, New York Dordrecht Heidelberg London, pp 1–48

Ron M, Kajala K, Pauluzzi G, Wang D, Reynoso MA, Zumstein K, Garcha J, Winte S, Masson H, Inagaki S, Federici F, Sinha N, Deal RB, Bailey-Serres J, Brady SM (2014) Hairy root transformation using *Agrobacterium rhizogenes* as a tool for exploring cell type-specific gene expression and function using tomato as a model. Plant Physiol 166: 455–469

Shimizu K, Aoki K (2019) Development of Parasitic Organs of a Stem Holoparasitic Plant in Genus *Cuscuta*. Front Plant Sci 10: 1435

Sun G, Xu Y, Liu H, Sun T, Zhang J, Hettenhausen C, Shen G, Qi J, Qin Y, Li J, Wang L, Chang W, Guo Z, Baldwin IT, Wu J (2018) Large-scale gene losses underlie the genome evolution of parasitic plant *Cuscuta australis*. Nat Commun 9: 2683

Svubova R, Blehova A (2013) Stable transformation and actin visualization in callus cultures of dodder (*Cuscuta europaea*). Biologia 68: 633–640

Tada Y, M. S, Furuhashi K (1996) Haustoria of *Cuscuta japonica*, a holoparasitic flowering plant, are induced by the cooperative effects of raf-red light and tactile stimuli. Plant Cell Physiol 37: 1049–1053

Vaughn KC (2002) Attachment of the parasitic weed dodder to the host. Protoplasma 219: 227–237

Vogel A, Schwacke R, Denton AK, Usadel B, Hollmann J, Fischer K, Bolger A, Schmidt MH, Bolger ME, Gundlach H, Mayer KFX, Weiss-Schneeweiss H, Temsch EM, Krause K (2018) Footprints of parasitism in the genome of the parasitic flowering plant *Cuscuta campestris*. Nat Commun 9: 2515

Yuncker TG (1932) The genus *Cuscuta*. Memoirs of the Torrey Botanical Club 18: 113–331

Zhang Y, Wang D, Wang Y, Dong H, Yuan Y, Yang W, Lai D, Zhang M, Jiang L, Li Z (2020) Parasitic plant dodder (*Cuscuta* spp.): A new natural *Agrobacterium*-to-plant horizontal gene transfer species. Sci China Life Sci 63: 312–316

